# The integrated stress response induces a common cell-autonomous death receptor 5-dependent apoptosis switch

**DOI:** 10.1101/2022.07.04.498696

**Authors:** Nerea L. Muniozguren, Francesca Zappa, Diego Acosta-Alvear

## Abstract

The integrated stress response (ISR) is a fundamental signaling network that reprograms the transcriptome and proteome to leverage the cell’s biosynthetic capacity against different stresses. Signaling plasticity is enabled by distinct ISR sensor kinases that detect specific perturbations. The ISR is dichotomous, with tailored homeostatic outputs and a terminal one engaged upon overwhelming stress. Through a chemical-genetics approach that uncouples natural stress inputs from ISR actuation, we show that the ISR engages an input-agnostic, cell-autonomous apoptosis mechanism that requires unconventional signaling by death receptor 5. Our results indicate that a common ISR mechanism eliminates terminally injured cells.

## Introduction

The integrated stress response (ISR) is an evolutionary conserved intracellular signaling network governed by four sensor kinases—GCN2, HRI, PKR, and PERK—activated by specific stress inputs (Reviewed in Costa-Mattioli & Walter, 2020). GCN2 detects uncharged amino acids and ribosome collisions (Wu *et al*, 2020; Dong *et al*, 2000), HRI detects heme deficits and relays mitochondrial stress (Han *et al*, 2001; Guo *et al*, 2020), PKR detects viral and endogenous double-stranded RNAs (dsRNAs) (Ehrenfeld & Hunt, 1971; Kim *et al*, 2014, 2018; Elbarbary *et al*, 2013), and PERK detects loss of ER proteostasis (Wang *et al*, 2018; Harding *et al*, 2000). All ISR kinases converge on phosphorylating a single serine in the alpha subunit of the eukaryotic initiation factor 2 (eIF2α), a heterotrimeric GTPase that, together with GTP and the initiator methionyl tRNA, forms the ternary complex which is required to initiate translation. Phosphorylated eIF2α acts as a competitive inhibitor of its guanine nucleotide exchange factor eIF2B (Adomavicius *et al*, 2019; Schoof *et al*, 2021), and as such, it decreases the availability of ternary complex leading to a global shutdown of protein synthesis. However, some mRNAs harboring upstream regulatory open reading frames (uORFs) escape this regulatory control and are selectively translated upon eIF2α phosphorylation. These mRNAs include those encoding the transcription factors ATF4 and CHOP, as well as *GADD34*, which encodes a regulatory subunit of protein phosphatase 1 that dephosphorylates eIF2α and establishes a negative feedback loop that terminates ISR signaling (Novoa *et al*, 2003; Vattem & Wek, 2004; Hinnebusch *et al*, 2016).

Through activation of gene expression programs and translational control, the ISR reprograms the transcriptome and proteome to restore homeostasis. However, during persistent stress, the ISR can switch to drive apoptosis. Both outcomes, adaptation and cell death, protect the organism by preserving healthy cells and eliminating terminally damaged ones. Signaling dynamics and intercommunication between nodes could explain the dichotomy between an adaptive and a terminal ISR. Indeed, during ER stress, opposing signals between PERK and the evolutionarily conserved ER stress sensor kinase/RNase IRE1 dictate adaptive or terminal outcomes (Lu *et al*, 2014). In response to ER stress, PERK induces CHOP, which in turn induces Death Receptor 5 (DR5), a transmembrane receptor belonging to the Tumor Necrosis Factor (TNF) superfamily of death receptors together with DR4, TNF, and Fas (Yamaguchi & Wang, 2004). Ligand binding promotes self-association of DR5 and recruitment of the adaptor protein FADD and procaspase-8 to nucleate the Death-Inducing Signaling Complex (DISC), which processes procaspase-8 into active caspase-8 (Wilson *et al*, 2009). In the early, adaptive phase of the ER stress response, IRE1 cleaves the DR5 mRNA, which leads to its degradation. Dampening of pro-survival IRE1 signaling during persistent ER stress switches the balance in favor of pro-apoptotic signaling downstream of the PERK-CHOP axis, which results in accumulation and activation of DR5. Notably, in this mechanism, DR5 accumulates in the *cis*-Golgi apparatus and signals unconventionally, without the need for its cognate ligand TRAIL, to engage a cell-autonomous death program (Lu *et al*, 2014; Lam *et al*, 2020).

Prompted by these observations, we wondered whether the ISR, independently of ER stress, engages a universal cell-autonomous apoptosis program dependent on unconventional DR5 signaling. Here we show that DR5 is induced by different ISR stress inputs, indicating a common ISR cell-death switch. Moreover, stress-free activation of the ISR using a chemical-genetics approach led to the accumulation of DR5 in the *cis*-Golgi apparatus and ligandin-dependent signaling reminiscent of that occurring during persistent ER stress. Our results suggest that the terminal ISR operates through a stress input-agnostic, cell-autonomous, universal apoptosis mechanism.

## Results

### Different ISR stress inputs induce DR5 and apoptosis

To dissect whether terminal ISR signals downstream of different stress inputs converge on DR5 expression, we treated H4 neuroglioma cells with various stressors that activate each of the ISR kinases. We chose a neural cell line because a dysregulated ISR has been observed in numerous neuropathologies (Costa-Mattioli & Walter, 2020). To induce ER stress and activate PERK, we treated cells with the ER calcium reuptake inhibitor thapsigargin (Schröder, 2008; Oslowski & Urano, 2011). To induce dsRNA stress and activate PKR, we transfected cells with the dsRNA mimetic polyinosinic-polycytidylic acid (poly I:C) (Balachandran *et al*, 2000). To induce mitochondrial stress and activate HRI, we treated cells with the ATP synthase inhibitor oligomycin (Guo *et al*, 2020). Finally, to mimic nutritional deficit and activate GCN2, we treated cells with histidinol, a histidine analog alcohol that prevents histidyl-tRNA charging (Harding *et al*, 2019). We next measured the levels of DR5 mRNA by qRT-PCR upon exposure of the cells to the abovementioned stressors for 18 hours, which we reasoned would be sufficient to initiate a terminal response based on previous observations (Lu *et al*, 2014) (Fig. 1A). This analysis revealed an approximately 4-fold upregulation of the DR5 mRNA in response to ER stress elicited by thapsigargin (Fig. 1A), consistent with the upregulation of DR5 mRNA observed in colon cancer cells subjected to persistent ER stress (Yamaguchi & Wang, 2004). Poly I:C, oligomycin, and histidinol also elevated the levels of the DR5 mRNA, albeit less potently, with approximately 2-fold (poly I:C and oligomycin) to 3-fold (histidinol) increases. Notably, we did not detect the mRNA encoding the death receptor DR4, which encodes a protein related to DR5 (Wilson *et al*, 2009) (Fig.S1A).

**Figure 1.**
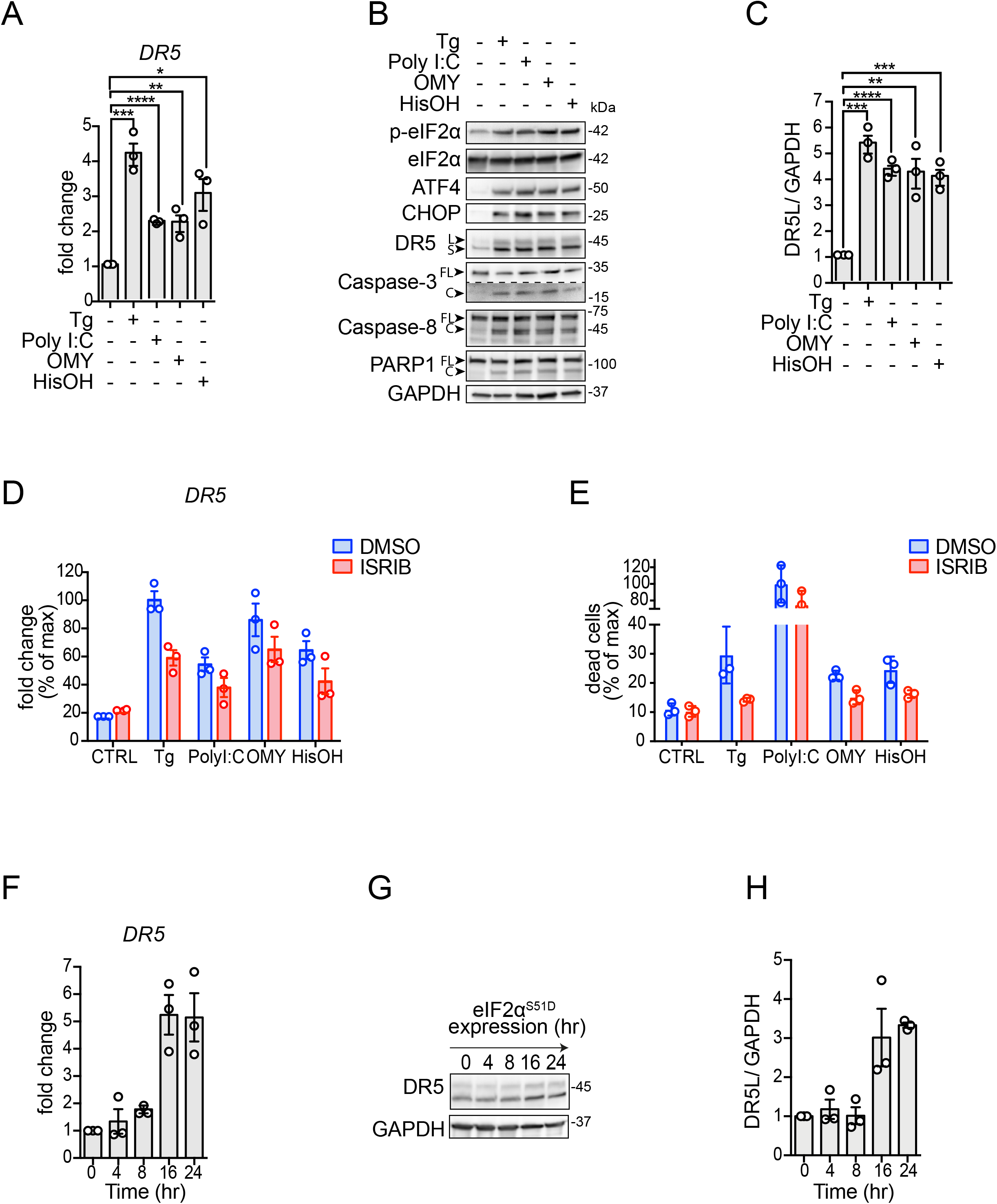
Multiple ISR inputs induce DR5-driven cell death. (A) Quantitative real-time PCR analysis of DR5 mRNA levels in H4 cells after activation of different branches of the ISR. Thapsigargin (Tg) 300 nM, poly I:C 250 ng/ml, oligomycin (OMY) 3 μM, histidinol (HisOH) 5 mM (mean and SEM, N = 3, \*\*\*\**P* < 0.0001, \*\*\**P* < 0.001, \*\**P* < 0.01, \**P* < 0.05 unpaired Student’s t-test, non-parametric). (B) Western blot showing upregulation of DR5, cleavage of caspase-8, caspase-3, and PARP1, and induction of canonical ISR markers (p-eIF2α, ATF4 and CHOP) in H4 cells upon treatment with different ISR stressors for 18 hours. GAPDH: loading control. (C) Densitometry quantification of the Western blot data for DR5 long isoforms (mean and SEM, N = 3, \*\*\*\**P* < 0.0001, \*\*\**P* < 0.001, \*\**P* < 0.01 unpaired Student’s t-test, non-parametric). (D) qRT-PCR analysis of DR5 mRNA levels after pre-treating H4 cells with the ISR inhibitor ISRIB (mean and SEM, N = 3, \*\*\*\**P* < 0.0001 One-way ANOVA). (E) Quantification of cell viability analysis using flow cytometry after staining with propidium iodide. The plot shows the induction of cell death in H4 cells upon treatment with classical pharmacological ISR activators and the restoration of cell viability upon ISR inhibition using ISRIB. Data are expressed as a percentage of the maximum value (mean and SEM, N = 3, \*\*\*\**P* < 0.0001 One-way ANOVA). (F) qRT-PCR analysis of DR5 mRNA levels in H4 cells expressing eIF2α ^S51D^ (mean and SEM, N = 3, \*\*\**P* < 0.001 One-way ANOVA). (G) Western blot showing upregulation of DR5 isoforms in H4 cells after activation of eIF2α^S51D^. GAPDH: loading control. (H) Densitometry quantification of the Western blot data for DR5 long isoform. GAPDH: loading control (mean and SEM, N = 3, \*\*\**P* < 0.001 One-way ANOVA).

The increase in DR5 mRNA elicited by any of the stressors we used was reflected at the protein level, with induction of both the long (DR5L) and short (DR5S) isoforms (DR5L, ranging from approximately 3.5-to 4.5-fold upregulation) (Valley *et al*, 2012) (Fig. 1B,C). The stress-induced upregulation of DR5 isoforms was accompanied by the processing of procaspase-8, activation of caspase-3, and cleavage of the canonical apoptosis marker PARP1 (Fig. 1B), which is consistent with DR5’s pro-apoptotic activity. Expectedly, these changes tracked with activation of the ISR, as indicated by phosphorylation of the ISR sensor kinases and eIF2α, as well as induction of ATF4 and CHOP (Fig. 1B; S1B).

Given that DR5 is induced by CHOP (Yamaguchi & Wang, 2004), our results suggest that the cell death decision is relayed to terminal effectors by eIF2α phosphorylation—the ISR core. If this is the case, inhibition of the ISR should suppress DR5 accumulation and apoptosis. To test this possibility, we co-treated cells with the small molecule ISR inhibitor ISRIB, which renders cells insensitive to the effects of eIF2α phosphorylation (Sidrauski *et al*, 2015) and the different ISR stressors aforementioned. ISRIB inhibited the upregulation of DR5 mRNA (Fig. 1D), and it restored cell viability, measured by the exclusion of the cell-impermeable DNA dye propidium iodide (PI) in live cells (Fig. 1E). These results indicate that cell-death signals can be bypassed in cells that are refractory to the effects of phosphorylated eIF2α and substantiate the notion that the ISR core passes the cell death decision to terminal effectors.

We next examined the ability of phosphorylated eIF2α to induce DR5 without any of the stress-inducing agents we used. To this end, we employed a genetics-based approach in which we force-expressed eIF2α^S51D^, a phosphomimetic point mutant of eIF2α, under the control of a tetracycline-regulatable promoter. Induction of eIF2α^S51D^ in H4 cells treated with doxycycline revealed a time-dependent accumulation of DR5 mRNA, starting at 8 hours after induction and reaching saturation at 16 hours (Fig. 1F). This time frame is consistent with the expression of the eIF2α^S51D^ and consequent activation of the ISR (Fig. S1C-E). Furthermore, the upregulation of DR5 mRNA was mirrored by DR5L and DR5S proteins (average 3-fold induction at 16 hours for DR5L) (Figs. 1G, H). Together, these results indicate that DR5 is induced by the different branches of the ISR downstream of phosphorylated eIF2α.

### Stress-free activation of the ISR induces DR5 and apoptosis

To dissect the molecular circuitry exclusive to the terminal ISR and avoid the pleiotropic effects of stress-inducing agents, we employed a chemical-genetics approach consisting of an engineered ISR sensor kinase, FKBP-PKR, which can be activated with a small molecule ligand to induce the ISR (Zappa *et al*, 2022). Unlike the classical ISR stress-inducing agents mentioned above, FKBP-PKR allows a precise, stress-free activation of a “pure” ISR. Treatment of cells bearing FKBP-PKR with the small molecule ligand led to a greater than 4-fold induction of the DR5 mRNA, with levels that peaked at 8 hours after adding the small molecule ligand (Fig. 2A).

**Figure 2.**
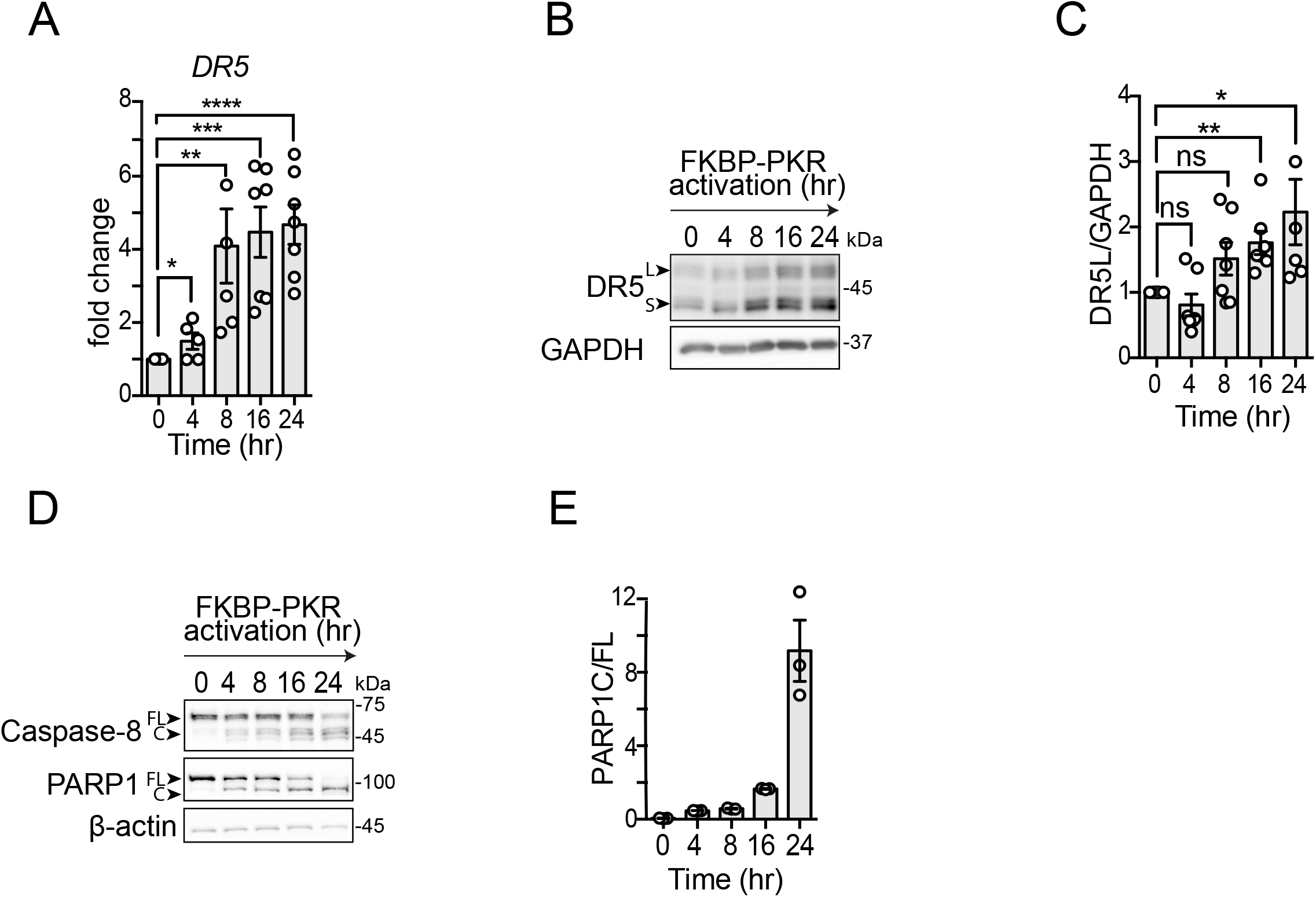
Stress-free activation of the ISR induces DR5 and caspase-8 cleavage. (A) qRT-PCR analysis of DR5 mRNA levels after activation of FKBP-PKR in H4 cells (mean and SEM, N = 7, \*\*\*\**P* < 0.0001, \*\*\**P* < 0.001, \*\**P* < 0.01, \**P* < 0.05 unpaired Student’s t-test, non-parametric). (B) Western blot showing upregulation of DR5 isoforms after activation of FKBP-PKR. GAPDH: loading control. (C) densitometry quantification of the long DR5 isoform (mean and SEM, N = 5, \*\**P* < 0.01, \**P* < 0.05, ns= not significant, unpaired Student’s t-test, non-parametric) (D) Western blot showing caspase-8 and PARP1 cleavage after FKBP-PKR activation. GAPDH: loading control. (E) Densitometry quantification of PARP1 cleavage upon FKBP-PKR activation at the indicated time points (mean and SEM, N=3).

The rise in DR5 mRNA levels was mirrored by a time-dependent accumulation of DR5 (approximate 2-fold induction at 16 hours for DR5L) protein after the addition of the small molecule ligand (Fig. 2B,C). Expectedly, these changes in DR5 were accompanied by the processing of procaspase-8 and cleavage of PARP1 upon activation of FKBP-PKR with the small molecule ligand (Fig. 2D,E). Together, these results indicate that DR5 can be induced by CHOP in a stress-input agnostic manner to initiate cell-autonomous apoptosis downstream of the ISR.

### Apoptosis downstream of the ISR requires DR5

To test the dependence of the ISR cell death program on DR5, we knocked down DR5 using CRISPR interference (CRISPRi) in H4 cells in cells bearing FKBP-PKR (Fig. S3A) and monitored the induction of apoptosis upon treatment with the small molecule ligand. CRISPRi-mediated depletion of DR5 resulted in a significant reduction in the activation of caspase-8 and caspase-3 (Fig. 3A), substantiating the notion that DR5 is a primary determinant of ISR-induced apoptosis.

**Figure 3.**
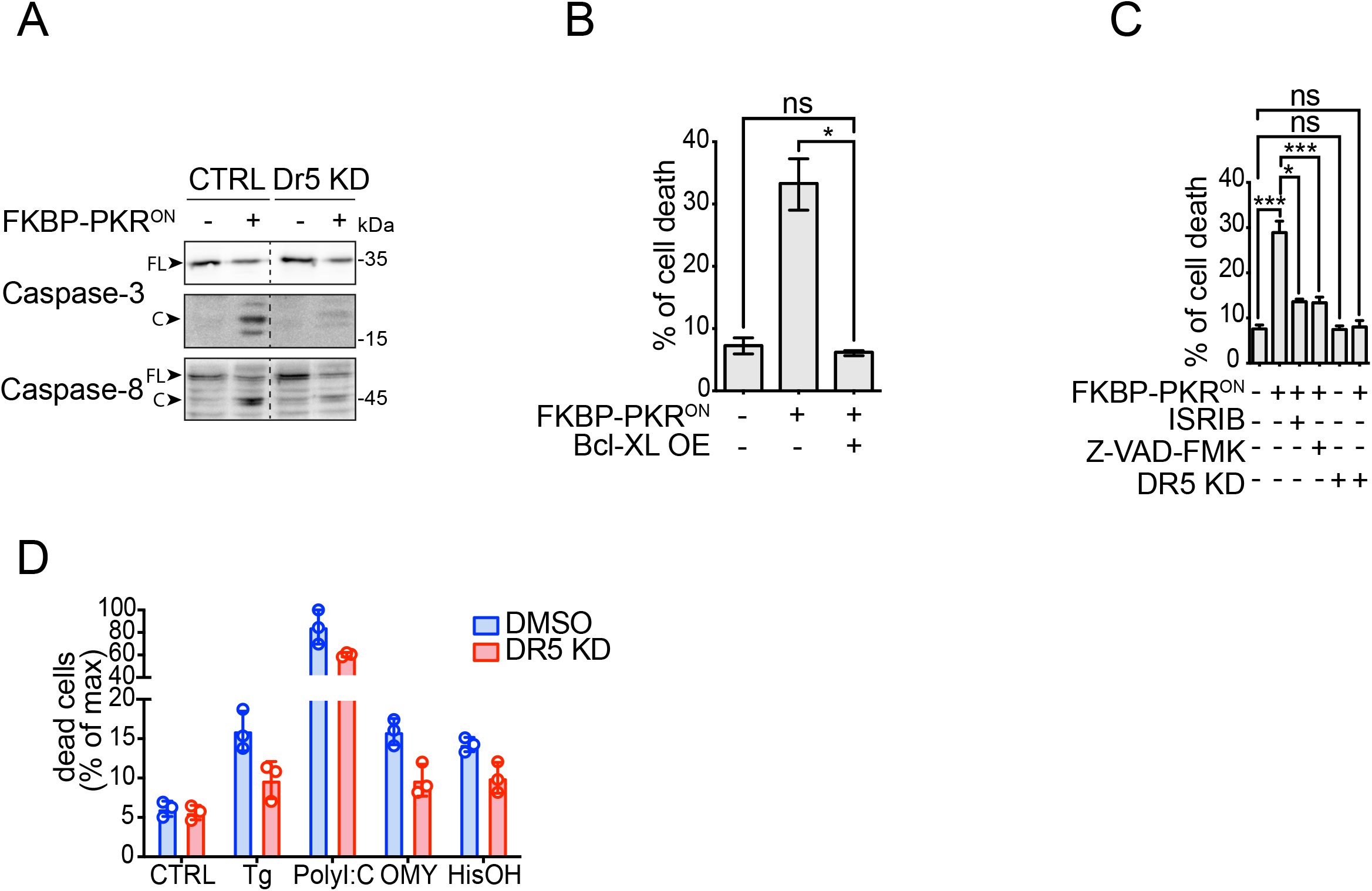
Apoptosis downstream of the ISR requires DR5 and caspase activity. (A) Western blot showing a lack of caspase-8 and caspase-3 activation upon DR5 knockdown in H4 cells in which we activated FKBP-PKR. (B) Flow cytometry quantification of cell death after propidium iodide staining of H4 cells in which we activated FKBP-PKR and overexpressing BCL-XL (mean and SEM, N = 3, \**P* < 0.05, unpaired Student’s t-test, non-parametric). (C) Flow cytometry quantification of cell death after propidium iodide staining in H4 cells treated with the ISR inhibitor ISRIB, the pan-caspase inhibitor Z-VAD-FMK, and upon genetic depletion of DR5 by CRISPRi (mean and SEM, N = 3, \*\*\**P* < 0.001, \**P* < 0.05, ns= not significant, unpaired Student’s t-test, non-parametric). (D) Flow cytometry quantification of cell death after propidium iodide staining in H4 DR5 CRISPRi cells treated with ISR pharmacological activators (mean and SEM, N = 3, \*\*\*\**P* < 0.0001 One-way ANOVA).

Apoptosis is controlled by extrinsic (death receptor-dependent) and intrinsic (mitochondria-dependent) interconnected signaling pathways that converge on the activation of executioner caspases. The pro-apoptotic protein BID, cleaved by caspase-8, bridges the extrinsic and intrinsic pathways (Fulda & Debatin, 2006). The active, truncated form of BID, tBID, promotes mitochondrial membrane permeabilization and cytochrome c release, triggering apoptosome formation and caspase-9 activation (Korsmeyer *et al*, 2000). Thus, we reasoned that a stress-free ISR induced by activation of FKBP-PKR enlists the intrinsic apoptosis pathway downstream of caspase-8. To test whether the ISR engages intrinsic apoptosis signals, we stably overexpressed the pro-survival protein BCL-XL, which inhibits mitochondrial membrane permeabilization (Billen *et al*, 2008) in cells bearing FKBP-PKR treated with the small molecule ligand. These experiments indicated that forced expression of BCL-XL forestalled cell death elicited by activation of FKBP-PKR as measured by PI staining (Fig. 3B).

Expectedly, and attesting to ISR involvement, treatment of cells in which we activated FKBP-PKR with ISRIB restored cell viability almost completely, as did treatment with the pancaspase inhibitor Z-VAD-FMK (approximately 28% cell death down to 12% Fig. 3C compare columns 1 to columns 3 and 4, respectively), corroborating ISR and caspase involvement (Fig. 3C). Notably, the genetic depletion of DR5 fully restored cell viability in cells in which we activated FKBP-PKR to levels that mirrored those of the untreated controls (approximately 28% cell death down to 8% Fig. 3C compare columns 1 and 6), which is consistent with a potential incomplete penetrance of the drug treatments when compared to genetic manipulation. Moreover, the depletion of DR5 alone had no effects on cell viability (Fig. 3C compare columns 1 and 5), and knockdown of DR5 strongly restored cell viability in H4 cells treated with pharmacological ISR inducers (Fig. 3D, S3C), further substantiating the notion that DR5 is required to induce apoptosis in the terminal ISR.

### Stress-free activation of the ISR leads to intracellular ligand-independent activation of DR5

As mentioned before, during persistent ER stress—which activates the PERK branch of the ISR— DR5 activates intracellularly independently of its ligand TRAIL (Lu *et al*, 2014; Lam *et al*, 2020). Based on this observation, we wondered whether DR5 accumulates intracellularly and signals similarly upon induction of a stress-free ISR. To answer this question, we first measured the levels of TRAIL mRNA by qRT-PCR and found that TRAIL mRNA levels rise in cells bearing FKBP-PKR in response to treatment with the small molecule ligand. Still, its upregulation subsides in a time-dependent manner (Fig. 4A, note that the qRT-PCR cycle threshold value (Ct) first decreases at 4 hours after FKBP-PKR activation and increases as time advances, indicating a rise and decay of the TRAIL mRNA levels). These results suggest that TRAIL might be induced in cells in which we activate the ISR, albeit at modest levels.

**Figure 4.**
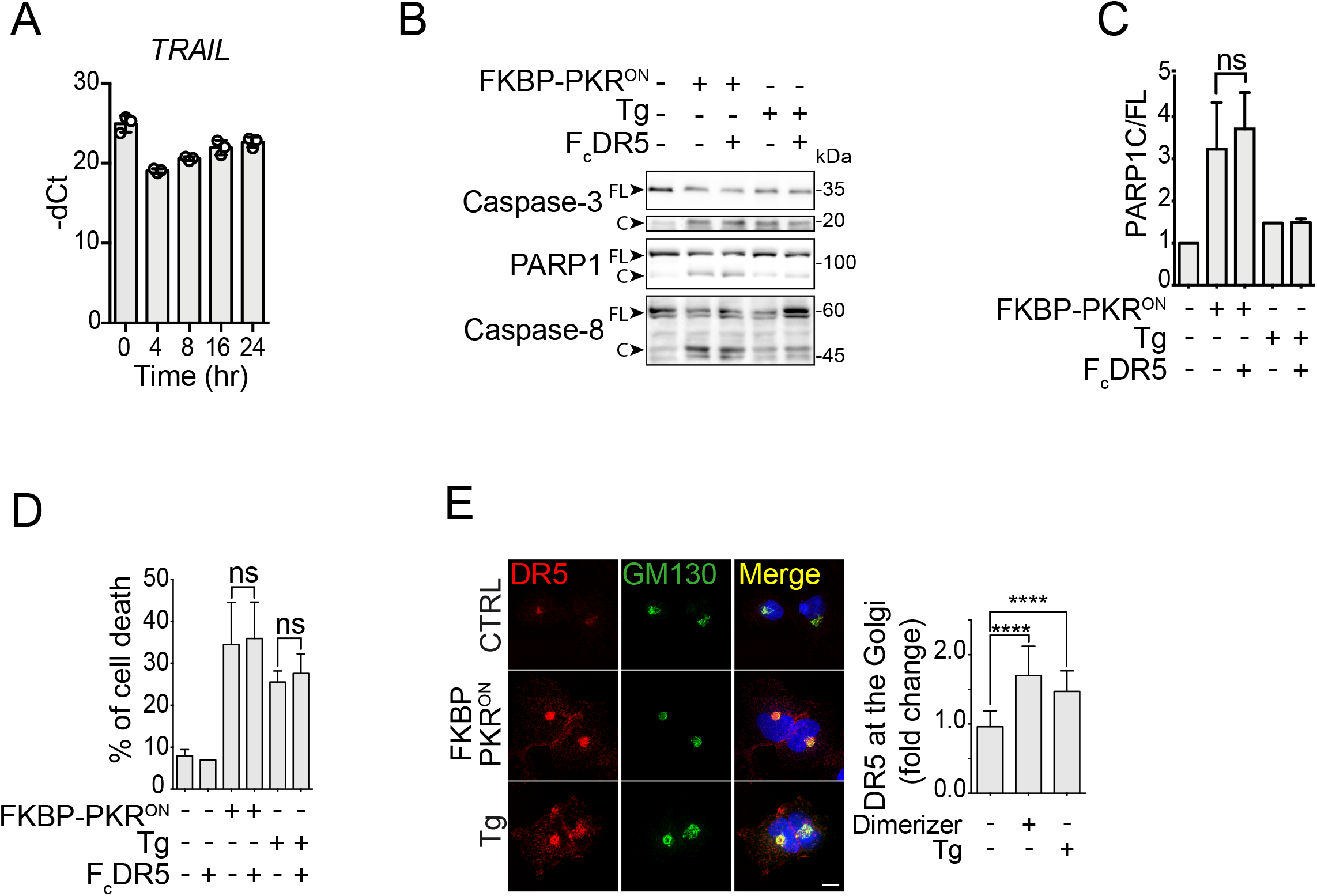
Stress-free activation of the ISR leads to intracellular ligand-independent activation of DR5. (A) Quantitative real-time PCR analysis showing the levels of TRAIL mRNA after activation of FKBP-PKR. (B) Western blot showing that blocking plasma membrane DR5 with FcDR5 does not impede the activation of caspase-3, caspase-8, and PARP1 cleavage in H4 cells in which we activated FKBP-PKR. (C) densitometry quantification of PARP1 cleavage upon activation of FKBP-PKR and co-treatment with FcDR5 (mean and SEM, N = 3, ns = not significant, unpaired Student’s t-test, non-parametric). (D) Flow cytometry quantification of cell death after propidium iodide staining of H4 cells treated with FcDR5 or thapsigargin (Tg, positive control). (E) Representative immunofluorescence images showing that DR5 co-localizes with the *cis*-Golgi apparatus marker GM130 upon induction of the ISR with natural stress inputs or in stress-free conditions after. activation of FKBP-PKR. Right panel: quantification of extent of localization of DR5 in the *cis*-Golgi apparatus in immunofluorescence analyses. (mean and SEM, N = 3, n>1000, \*\*\*\**P* < 0.001, \**P* < 0.05, unpaired Student’s t-test, non-parametric).

The observation that TRAIL might be induced during the ISR suggests it could engage DR5 at the plasma membrane. To test whether plasma membrane-localized DR5 signals apoptosis, we exposed cells in which we activated FKBP-PKR with the small molecule ligand to a DR5-neutralizing Fc antibody fragment (FcDR5). Treatment with FcDR5 did not prevent the activation of caspases-3 and −8, or cleavage or PARP1 (Fig. 4B,C), nor did it block cell death (Fig. 4D) in response to FKBP-PKR activation, indicating that plasma membrane DR5 is not required for transducing death signals upon activation of a terminal ISR.

Finally, we examined the subcellular localization of DR5 upon induction of a stress-free ISR by activation of FKBP-PKR. Treatment of cells bearing FKBP-PKR with the small molecule ligand led to an accumulation of DR5 in the *cis*-Golgi apparatus, as evidenced by co-staining with the *cis*-Golgi apparatus marker GM130 in immunofluorescence analyses (Fig. 4E). Strikingly, the intracellular localization of DR5 to the *cis*-Golgi apparatus elicited by a stress-free activation of the ISR was virtually indistinguishable from that caused by ER stress-inducing agents (Fig. 4E and (Lu *et al*, 2014; Lam *et al*, 2020), indicating that accumulation of DR5 in the *cis*-Golgi apparatus is a common feature of the terminal ISR. Together, these results substantiate that the terminal ISR engages a common, unconventional, cell-autonomous apoptosis mechanism that relies on TRAIL-independent intracellular DR5 activation.

## Discussion

Using orthogonal approaches, we demonstrate that the terminal ISR appoints a cell-autonomous apoptosis mechanism that relies on intracellular activation of DR5 and engagement of the extrinsic and intrinsic apoptosis pathways. We base this conclusion on multiple lines of evidence. First, natural lethal ISR stress inputs upregulate DR5 mRNA and protein (Figs. 1A, B), and actuate apoptosis downstream of DR5 (Fig. 1B), as does the stress-free activation of the ISR using a synthetic biology tool (Figs. 2A-E). Second, the ISR inhibitor ISRIB reverses the upregulation of DR5 and subsequent cell death triggered by the ISR (Fig. 1C, 3C), indicating that cell death signals pass through phosphorylation of eIF2α. Third, cells lacking DR5 are less susceptible to cell death triggered by classical ISR inducers (Fig. 3D) as well as by a stress-free ISR (Fig. 3C). Fourth, DR5 accumulates in the *cis*-Golgi apparatus in response to classical ISR inducers and upon stress-free induction of the ISR (Fig. 4E) and blocking plasma-membrane DR5 with a neutralizing antibody had no effect on cell viability (Fig. 4B, 4C), indicating intracellular, ligand-independent DR5 activation during the terminal ISR.

For cell health to ensue, the ISR must accurately interpret information about stress states and actuate accordingly to control homeostatic or terminal outputs. On the one hand, tailored homeostatic outcomes are likely executed through ISR kinase signal codes, which could result from additional ISR kinase substrates and interactors and from accessibility to different pools of eIF2α. On the other hand, considering all ISR sensor kinases pass signals through a core relay, eIF2α, the terminal ISR is likely to employ an off-the-shelf mechanism downstream of eIF2α that funnels information about an irreparable critical state to a common executor, DR5. Our data support this notion, indicating that the terminal ISR operates through a pro-apoptotic relay consisting of phospho-eIF2α→CHOP→DR5 (Yamaguchi & Wang, 2004; Lu *et al*, 2014).(Castelli *et al*, 1998; Besch *et al*, 2009; Siddiqui *et al*, 2015).

It is noteworthy that during ER stress, intracellular DR5 has been postulated to be activated by unfolded protein ligands (Lam *et al*, 2020). While we cannot formally exclude the possibility that our synthetic ISR activation approach induces some mild accumulation of unfolded proteins that can serve as DR5 activating ligands, it is unlikely it does so to a level that is comparable to that elicited by classic ER poisons (e.g., thapsigargin or the N-linked glycosylation inhibitor tunicamycin) or forced-expression of a single ER folding-mutant protein, such as myelin protein zero (Lam *et al*, 2020). Nevertheless, considering that some misfolded proteins bind DR5, it remains plausible that a sustained ISR may induce the accumulation of a single or a subset of select ER client protein(s) that may engage DR5 leading to its activation. Alternatively, DR5 crowding in the Golgi apparatus may be sufficient to initiate signaling. Indeed, DR5 overexpression triggers apoptosis (Screaton *et al*, 1997). Future -omics studies (coupled RNAseq and proteomics) in cells in which we activate the ISR synthetically may shed light on the mechanism of activation of DR5 during the terminal ISR. It is intriguing that the terminal ISR engages a cell surface death receptor unconventionally. Because plasma membrane DR5 is not required for the terminal ISR, and the bulk of the protein accumulates in the Golgi apparatus during the ISR, additional mechanisms may retain DR5 in the secretory pathway. This is an intriguing possibility that remains to be investigated.

Our results also indicate a potential node for therapeutic intervention in the ISR. A dysregulated ISR has been observed in numerous neurocognitive disorders (Bando *et al*, 2005; Zhu *et al*, 2019; Bond *et al*, 2020; Krukowski *et al*, 2020; Halliday *et al*, 2015). Therefore, a terminal ISR may lead to neural cell loss. Antisense oligonucleotides, which have shown efficacy in neurodegenerative diseases including Duchenne’s muscular dystrophy, spinal muscular atrophy, and hereditary transthyretin-mediated amyloidosis (Nguyen & Yokota, 2019; Relizani & Goyenvalle, 2018; Chen, 2019; Wood *et al*, 2017; Gales, 2019), could be deployed to target DR5 and prevent neural cell loss. Regardless of potential therapeutic applications, identifying a common mechanism controlling cell death during prolonged ISR signaling advances our understanding of how the ISR operates to maintain the health of tissues: customized homeostatic solutions or a one-size-fits-all terminal response.

---

**Figure S1.**
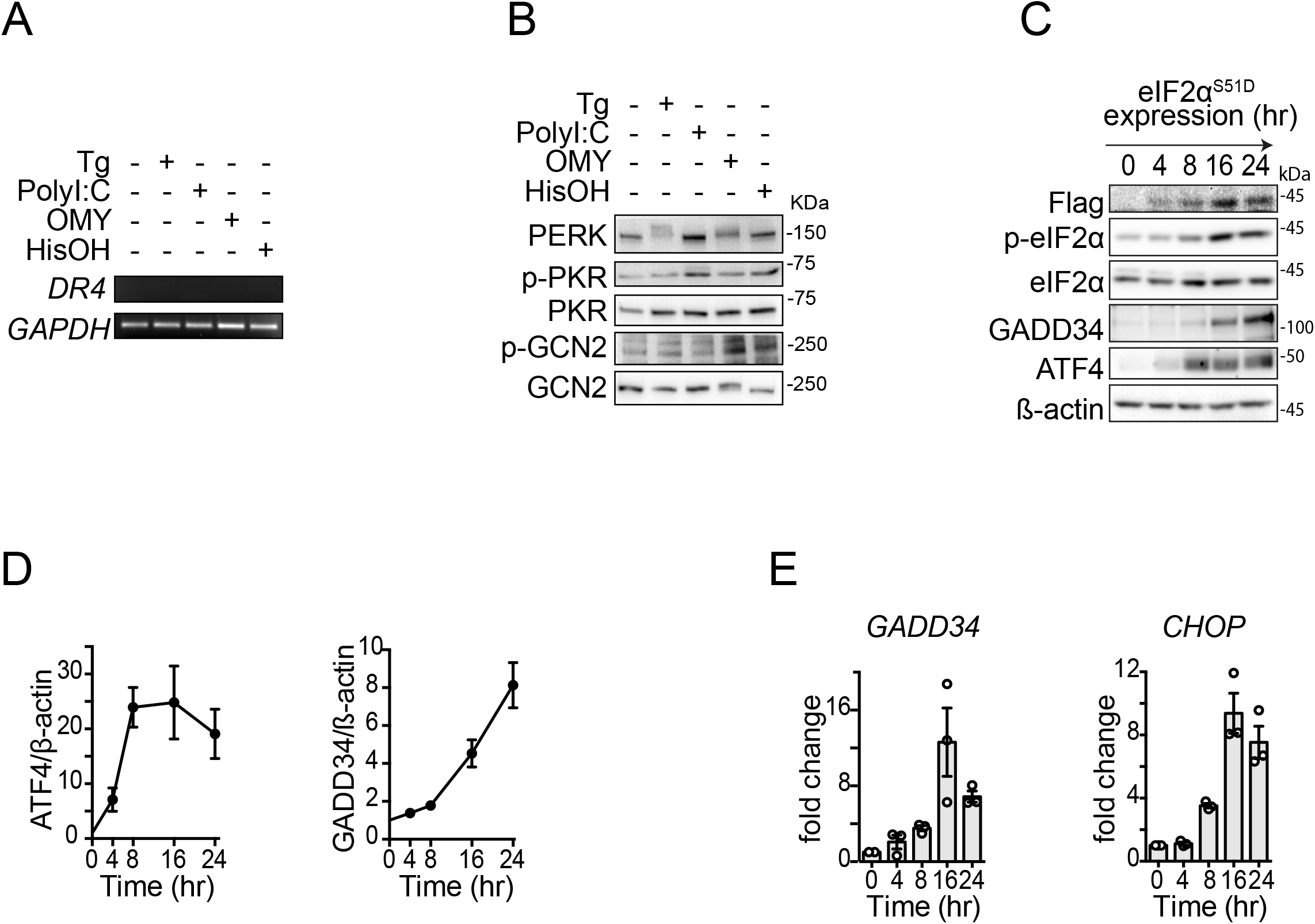
Pharmacological and genetic ISR induction in H4 cells. (A) Analysis of DR4 mRNA levels by RT-PCR in H4 cells after activation of the ISR with different pharmacological agents shows undetectable levels of DR4 transcript. GAPDH: loading control. Thapsigargin (Tg) 300 nM, poly I:C 250 ng/ml, oligomycin (OMY) 3 μM, histidinol (HisOH) 5 mM. (B) Western blot showing phosphorylation of the ISR sensor kinases in H4 cells upon treatment with pharmacological ISR inducers. (C) Western blot showing canonical ISR induction in H4 cells expressing FLAG epitope-tagged eIF2α^S51D^ and treated with doxycycline for the indicated time. β-actin: loading control. (D) Densitometry quantification of the Western blot data in (C) for ATF4 and GADD34 (mean and SEM, N = 3). (E) qRT-PCR analysis of ATF4 and CHOP mRNA levels in H4 cells expressing FLAG epitope-tagged eIF2α^S51D^ and treated with doxycycline for the indicated time (mean and SEM, N = 3).

**Figure S3.**
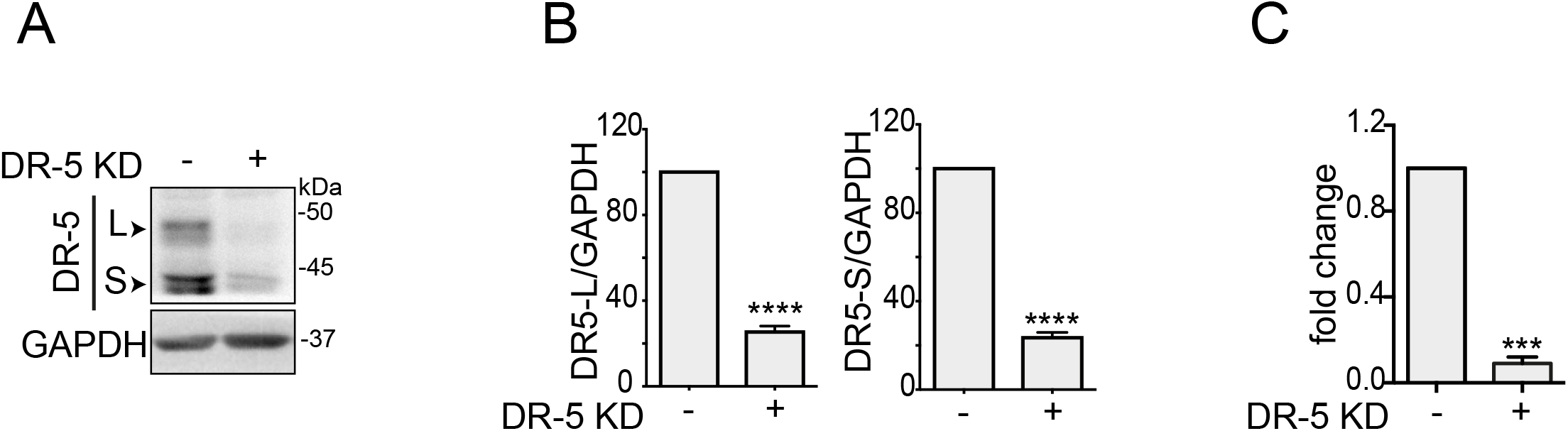
Generation of DR5-deficient cell lines. (A) Western blot showing the extent of CRISPRi-mediated knock-down of DR5 in H4 FKBP-PKR cells. GAPDH; loading control. (B) Densitometry quantification of the DR5 short and long isoforms upon genetic depletion by CRISPRi (mean and SEM, N = 3, \*\*\*\**P* < 0.0001, unpaired Student’s t-test, non-parametric). (C) Quantitative real-time PCR analysis showing the levels of DR5 in H4 cells (mean and SEM, N = 3, \*\*\**P* < 0.001, unpaired Student’s t-test, non-parametric).

## Materials and Methods

### Cell culture, genetic knockdown, transduction, and drug treatments

H4 cells expressing the CRISPRi machinery(Gilbert *et al*, 2014) were a kind gift from Martin Kampmann (UCSF). These cells were maintained in Dulbecco’s Modified Eagle Medium (DMEM) supplemented with 10% fetal bovine serum (FBS) and penicillin/streptomycin. DR5 gene was depleted using CRISPRi as previously described(Gilbert *et al*, 2014). Briefly, CRISPRi cells were transduced with VSV-G pseudotyped lentiviruses harboring the three top small guide RNAs (5’-GGGCAAGACGCACCAGTCGT-3’; 5’-GAGAGATGGGTCCCCGGGTT-3’; 5’-GAAAGTAGATCGGGCATCGT-3’). The sgRNA sequences were obtained from the human genome-scale CRISPRi library developed by the laboratory of Jonathan Weissman (MIT, Whitehead Institute).

For data collection, cells were seeded at a density of 1-2 × 10^5^ cells/well in 6-well plates, 0.7-1.0 × 10^5^ cells/well in 12-well plates, or 0.5-0.7 × 10^5^ cells/well in 24-well plates and maintained for a further 24 h before any treatment. Cells were treated with ISR stress inducers (300 nM thapsigargin (Sigma-Aldrich), 3 μM oligomycin (Sigma-Aldrich), 5 mM histidinol (Sigma-Aldrich), or transfected with 250 nM poly I:C (Tocris), as previously described(Zappa *et al*, 2022), 1μM ISRIB (Sigma-Aldrich), 50 μM Z-VAD-FMK (SelleckChem), or 1 μg/ml FcDR5 (R&D systems) as indicated. FKBP-PKR was activated with 100 nM of the homodimerization ligand AP20187 (Takara), as previously described(Zappa *et al*, 2022).

Stable cell lines bearing transgenes were generated by retroviral transduction as previously described(Sidrauski *et al*, 2013). Briefly, VSV-G pseudotyped retroviral particles encoding constructs of choice were prepared using standard protocols using GP2-293 packaging cells (Clontech). Viral supernatants were collected and used to infect target cells by centrifugal inoculation (spinoculation). For retroviral infections, target cells were plated at a density of 2 × 10^5^ cells/well in 6-well plates one day before transduction in presence of 8 μg/ml polybrene. Pseudoclonal cell populations were obtained by fluorescence-activated cell sorting using a narrow gate placed over the mean of the signal distribution as previously described (Zappa *et al*, 2022).

### Immunoblotting

Cell lysates were obtained by collecting cells directly in Laemmli SDS-PAGE sample buffer (62.5 mM Tris-HCl pH 6.8, 2% SDS, 10% glycerol and 0.01% bromophenol blue). Lysates were briefly sonicated and supplemented with fresh 5% 2-mercaptoethanol prior to heat denaturation and separation by SDS-PAGE. Lysates were separated 8-10% SDS-PAGE gels and transferred onto nitrocellulose membranes for immunoblotting. Immunoreactive bands were detected by enhanced chemiluminescence using horse radish peroxidase (HRP)-conjugated secondary antibodies. The antibodies and dilutions used were as follows: PKR (Cell Signaling Technology 3072, 1:2000), phospho-PKR (T466) (Abcam AB322036, 1:2000), eIF2α (Cell Signaling Technology 9722, 1:1000), phospho-eIF2α (Cell Signaling Technology 9721, 1:1000), FLAG (M2 clone Sigma Aldrich F1804, 1:2000), ATF4 (Cell Signaling Technology 11815, 1:1000), CHOP (Cell Signaling Technology 2895, 1:1000), anti-DR5 (Cell Signaling Technology 8074, 1:1000), PARP1(Cell Signaling 9532, 1:1000), caspase-8 (Cell Signaling Technology 9746, 1:1000), β-actin (1:5000, Sigma Aldrich, 061M4808), anti-GAPDH (Abcam 8245, 1:5000,), all diluted in 1% BSA-TBST. Secondary anti-rabbit and anti-mouse HRP-conjugated antibodies (Cell Signaling Technology 7074, 7076) were used at 1:5000 dilutions in 1% BSA-TBST.

### Immunofluorescence analyses

H4 cells were grown on coverslips in 24-well plates, fixed with 4% PFA for 10 min, washed three times with PBS, and permeabilized with blocking solution (0.05% saponin, 0.5% BSA, 50 mM, NH4Cl in PBS) for 20 min. DR5 (Cell Signaling Technology 8074, 1:200) and GM130 (BD technology 610822, 1:1000) primary antibodies were diluted in blocking solution and incubated for 1 hour at room temperature. The coverslips were washed with PBS and incubated with fluorochrome-conjugated secondary antibodies (Alexa fluor anti-mouse 647; Invitrogen A32728 and Alexa fluor anti-rabbit 568; Invitrogen A11011 diluted at 1:500 dilution in blocking solution) and DAPI (0.1 μg/mL) for 45 minutes at RT. Fixed cells were washed 2 times in PBS and one time in ddH_2_O and mounted on coverslips with Mowiol. Imaged were acquired using a resonant scanning confocal microscope (Leica SP8) equipped with a Plan Apochromat 60 × NA 1.2 oil immersion objective. Fluorescence microscopy images were processed with Fiji (ImageJ: National Institutes of Health) software. To determine the proportion of DR5 in the *cis*-Golgi complex, each cell in the field of view was cropped and a single ROI was drawn manually to quantify the total DR5 fluorescence signal. GM130 signal was used to calculate the DR5 signal in the *cis*-Golgi complex using the Fiji plug-in “create selection”. An average of 200 cells per time point was considered for each replicate. The data were expressed as the ratio between DR5 MFI in the cis-Golgi compartment over the total DR5 MFI. Statistical significance for differences between groups was calculated using the unpaired Student’s t-test with the GraphPad Prism software. All data reported as mean ± s.e.m.

### Cell viability assays

To measure cell viability by flow cytometry, we collected detached cells and adherent cells. Detached cells were first collected by centrifugation. Adherent cells were collected by trypsinization. Both cell populations were pooled and resuspended in PBS supplemented with 2% FBS and 0.1 mg/ml RNase. Subsequently, propidium iodide (PI, 1.5 μg/ml) was added to the cell suspension. The samples were incubated on ice for 10 min and separated in an Attune cytPix flow cytometer. Flow cytometry data was analyzed using FlowJo (TreeStar, USA). The proportions of live and dead cells as determined by PI staining were used as cell viability metrics.

### DR5 neutralizing antibody assay

Cells bearing FKBP-PKR cells were washed 2 times with PBS and either fresh cell culture medium (control) or fresh cell culture medium supplemented with FcDR5 neutralizing antibody (1 μg/ml) overnight. The following day, the homodimerization ligand AP20187 was added to the cells and the cells were incubated for 24 hours prior to collection for analysis by immunoblotting or flow cytometry.

### qRT-PCR Analysis

Total RNA from sub-confluent H4 cells was isolated using the RNeasy RNA purification kit (Qiagen), following the manufacturer’s recommendations. 1 μg to total RNA was reverse transcribed with SuperScript VILO (Invitrogen) the manufacturer’s recommendations. The resulting cDNA was used as a template for real-time qPCR using PowerUp SYBR Green Master Mix (Applied Biosystems), according to the manufactures’s protocol. GAPDH or β-ACTIN were used as normalizing controls to estimate fold-change in mRNA expression. Oligonucleotide primers used in this study are provided in Table S1.

**Table S1.**
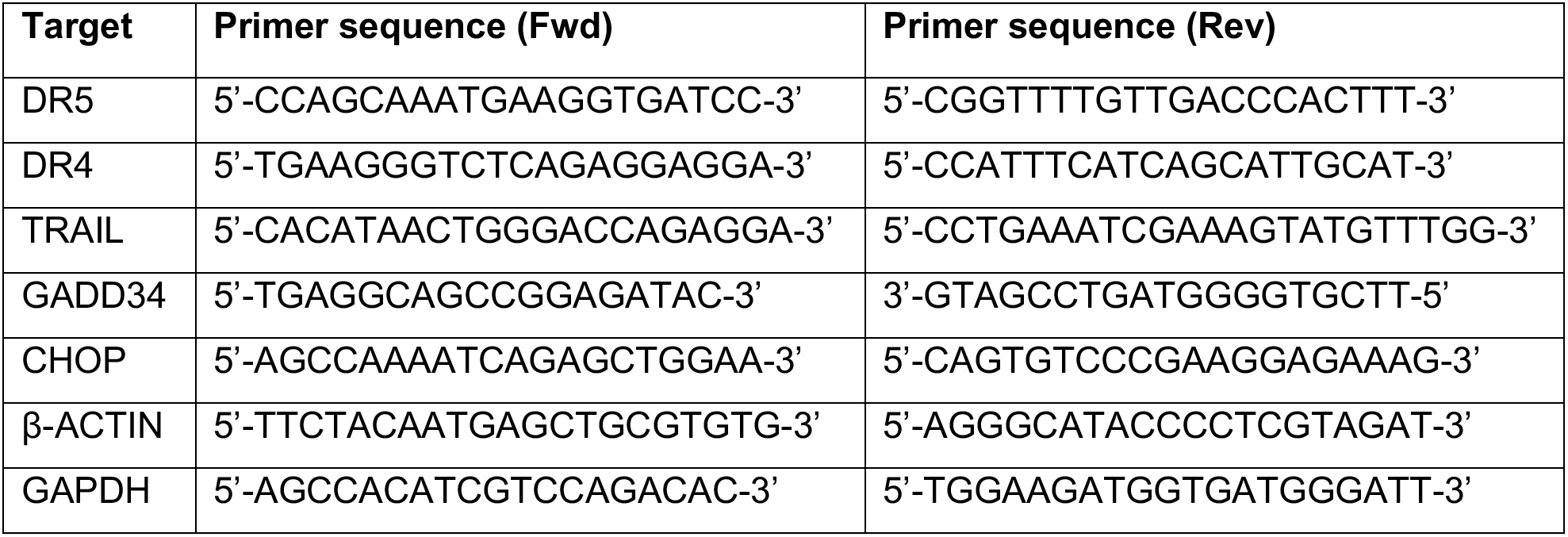

### Author contributions

D.A-A. supervised the research. N.L.M. and F.Z. performed experiments. F.Z. designed experiments and analyzed the data. N.L.M., D.A-A. and F.Z. wrote the manuscript.

### Author disclosures

D.A-A. is an inventor on U.S. patent 9708247 held by the Regents of the University of California that describes ISRIB and its analogs. Rights to the invention have been licensed to Calico Life Sciences LLC. For the rest of the authors, there are no competing interests.

## Bibliography

Adomavicius T, Guaita M, Zhou Y, Jennings MD, Latif Z, Roseman AM & Pavitt GD (2019) The structural basis of translational control by eIF2 phosphorylation. Nat Commun 10: 2136

Balachandran S, Roberts PC, Brown LE, Truong H, Pattnaik AK, Archer DR & Barber GN (2000) Essential Role for the dsRNA-Dependent Protein Kinase PKR in Innate Immunity to Viral Infection. Immunity 13: 129–141

Bando Y, Onuki R, Katayama T, Manabe T, Kudo T, Taira K & Tohyama M (2005) Doublestrand RNA dependent protein kinase (PKR) is involved in the extrastriatal degeneration in Parkinson’s disease and Huntington’s disease. Neurochem Int 46: 11–18

Besch R, Poeck H, Hohenauer T, Senft D, Häcker G, Berking C, Hornung V, Endres S, Ruzicka T, Rothenfusser S, et al (2009) Proapoptotic signaling induced by RIG-I and MDA-5 results in type I interferon–independent apoptosis in human melanoma cells. J Clin Invest 119: 2399–2411

Billen LP, Kokoski CL, Lovell JF, Leber B & Andrews DW (2008) Bcl-XL inhibits membrane permeabilization by competing with Bax. PLoS Biol 6: e147

Bond S, Lopez-Lloreda C, Gannon PJ, Akay-Espinoza C & Jordan-Sciutto KL (2020) The Integrated Stress Response and Phosphorylated Eukaryotic Initiation Factor 2α in Neurodegeneration. J Neuropathol Exp Neurol 79: 123–143

Castelli JC, Hassel BA, Maran A, Paranjape J, Hewitt JA, Li X, Hsu Y-T, Silverman RH & Youle RJ (1998) The role of 2′-5′ oligoadenylate-activated ribonuclease L in apoptosis. Cell Death Differ 5: 313–320

Chen I (2019) An antisense oligonucleotide splicing modulator to treat spinal muscular atrophy. Nature Research

Costa-Mattioli M & Walter P (2020) The integrated stress response: From mechanism to disease. Science 368: eaat5314

Dong J, Qiu H, Garcia-Barrio M, Anderson J & Hinnebusch AG (2000) Uncharged tRNA activates GCN2 by displacing the protein kinase moiety from a bipartite tRNA-binding domain. Mol Cell 6: 269–279

Ehrenfeld E & Hunt T (1971) Double-Stranded Poliovirus RNA Inhibits Initiation of Protein Synthesis by Reticulocyte Lysates. Proceedings of the National Academy of Sciences 68:1075–1078

Elbarbary RA, Li W, Tian B & Maquat LE (2013) STAU1 binding 3′ UTR IRAlus complements nuclear retention to protect cells from PKR-mediated translational shutdown. Genes Dev 27: 1495–1510

Fulda S & Debatin K-M (2006) Extrinsic versus intrinsic apoptosis pathways in anticancer chemotherapy. Oncogene 25: 4798–4811

Gales L (2019) Tegsedi (Inotersen): An Antisense Oligonucleotide Approved for the Treatment of Adult Patients with Hereditary Transthyretin Amyloidosis. Pharmaceuticals (Basel) 12: E78

Gilbert LA, Horlbeck MA, Adamson B, Villalta JE, Chen Y, Whitehead EH, Guimaraes C, Panning B, Ploegh HL, Bassik MC, et al (2014) Genome-Scale CRISPR-Mediated Control of Gene Repression and Activation. Cell 159: 647–661

Guo X, Aviles G, Liu Y, Tian R, Unger BA, Lin Y-HT, Wiita AP, Xu K, Correia MA & Kampmann M (2020) Mitochondrial stress is relayed to the cytosol by an OMA1-DELE1-HRI pathway. Nature 579: 427–432

Halliday M, Radford H, Sekine Y, Moreno J, Verity N, le Quesne J, Ortori CA, Barrett DA, Fromont C, Fischer PM, et al (2015) Partial restoration of protein synthesis rates by the small molecule ISRIB prevents neurodegeneration without pancreatic toxicity. Cell Death Dis 6: e1672–e1672

Han AP, Yu C, Lu L, Fujiwara Y, Browne C, Chin G, Fleming M, Leboulch P, Orkin SH & Chen JJ (2001) Heme-regulated eIF2alpha kinase (HRI) is required for translational regulation and survival of erythroid precursors in iron deficiency. EMBO J 20: 6909–6918

Harding HP, Ordonez A, Allen F, Parts L, Inglis AJ, Williams RL & Ron D (2019) The ribosomal P-stalk couples amino acid starvation to GCN2 activation in mammalian cells. eLife 8: e50149

Harding HP, Zhang Y, Bertolotti A, Zeng H & Ron D (2000) Perk Is Essential for Translational Regulation and Cell Survival during the Unfolded Protein Response. Molecular Cell 5: 897–904

Hinnebusch AG, Ivanov IP & Sonenberg N (2016) Translational control by 5’-untranslated regions of eukaryotic mRNAs. Science 352: 1413–1416

Kim Y, Lee JH, Park J-E, Cho J, Yi H & Kim VN (2014) PKR is activated by cellular dsRNAs during mitosis and acts as a mitotic regulator. Genes Dev 28: 1310–1322

Kim Y, Park J, Kim S, Kim M, Kang M-G, Kwak C, Kang M, Kim B, Rhee H-W & Kim VN (2018) PKR Senses Nuclear and Mitochondrial Signals by Interacting with Endogenous Double-Stranded RNAs. Mol Cell 71: 1051–1063.e6

Korsmeyer SJ, Wei MC, Saito M, Weiler S, Oh KJ & Schlesinger PH (2000) Pro-apoptotic cascade activates BID, which oligomerizes BAK or BAX into pores that result in the release of cytochrome c. Cell Death Differ 7: 1166–1173

Krukowski K, Nolan A, Frias ES, Boone M, Ureta G, Grue K, Paladini M-S, Elizarraras E, Delgado L, Bernales S, et al (2020) Small molecule cognitive enhancer reverses age-related memory decline in mice. eLife 9: e62048

Lam M, Marsters SA, Ashkenazi A & Walter P (2020) Misfolded proteins bind and activate death receptor 5 to trigger apoptosis during unresolved endoplasmic reticulum stress. Elife 9: e52291

Lu M, Lawrence DA, Marsters S, Acosta-Alvear D, Kimmig P, Mendez AS, Paton AW, Paton JC, Walter P & Ashkenazi A (2014) Opposing unfolded-protein-response signals converge on death receptor 5 to control apoptosis. Science 345: 98–101

Nguyen Q & Yokota T (2019) Antisense oligonucleotides for the treatment of cardiomyopathy in Duchenne muscular dystrophy. Am J Transl Res 11: 1202–1218

Novoa I, Zhang Y, Zeng H, Jungreis R, Harding HP & Ron D (2003) Stress-induced gene expression requires programmed recovery from translational repression. EMBO J 22: 1180–1187

Oslowski CM & Urano F (2011) Measuring ER stress and the unfolded protein response using mammalian tissue culture system. Methods Enzymol 490: 71–92

Relizani K & Goyenvalle A (2018) The Use of Antisense Oligonucleotides for the Treatment of Duchenne Muscular Dystrophy. Methods Mol Biol 1687: 171–183

Schoof M, Boone M, Wang L, Lawrence R, Frost A & Walter P (2021) eIF2B conformation and assembly state regulate the integrated stress response. eLife 10: e65703

Schröder M (2008) Endoplasmic reticulum stress responses. Cell Mol Life Sci 65: 862–894

Screaton GR, Mongkolsapaya J, Xu XN, Cowper AE, McMichael AJ & Bell JI (1997) TRICK2, a new alternatively spliced receptor that transduces the cytotoxic signal from TRAIL. Curr Biol 7: 693–696

Siddiqui MA, Mukherjee S, Manivannan P & Malathi K (2015) RNase L Cleavage Products Promote Switch from Autophagy to Apoptosis by Caspase-Mediated Cleavage of Beclin-1. Int J Mol Sci 16: 17611–17636

Sidrauski C, Acosta-Alvear D, Khoutorsky A, Vedantham P, Hearn BR, Li H, Gamache K, Gallagher CM, Ang KK-H, Wilson C, et al (2013) Pharmacological brake-release of mRNA translation enhances cognitive memory. eLife 2: e00498

Sidrauski C, McGeachy AM, Ingolia NT & Walter P (2015) The small molecule ISRIB reverses the effects of eIF2α phosphorylation on translation and stress granule assembly. eLife 4: e05033

Valley CC, Lewis AK, Mudaliar DJ, Perlmutter JD, Braun AR, Karim CB, Thomas DD, Brody JR & Sachs JN (2012) Tumor necrosis factor-related apoptosis-inducing ligand (TRAIL) induces death receptor 5 networks that are highly organized. J Biol Chem 287: 21265–21278

Vattem KM & Wek RC (2004) Reinitiation involving upstream ORFs regulates ATF4 mRNA translation in mammalian cells. Proc Natl Acad Sci U S A 101: 11269–11274

Wang P, Li J, Tao J & Sha B (2018) The luminal domain of the ER stress sensor protein PERK binds misfolded proteins and thereby triggers PERK oligomerization. J Biol Chem 293: 4110–4121

Wilson NS, Dixit V & Ashkenazi A (2009) Death receptor signal transducers: nodes of coordination in immune signaling networks. Nat Immunol 10: 348–355

Wood MJA, Talbot K & Bowerman M (2017) Spinal muscular atrophy: antisense oligonucleotide therapy opens the door to an integrated therapeutic landscape. Hum Mol Genet 26: R151–R159

Wu CC-C, Peterson A, Zinshteyn B, Regot S & Green R (2020) Ribosome Collisions Trigger General Stress Responses to Regulate Cell Fate. Cell 182: 404–416.e14

Yamaguchi H & Wang H-G (2004) CHOP is involved in endoplasmic reticulum stress-induced apoptosis by enhancing DR5 expression in human carcinoma cells. J Biol Chem 279: 45495–45502

Zappa F, Muniozguren NL, Wilson MZ, Costello MS, Ponce-Rojas JC & Acosta-Alvear D (2022) Signaling by the integrated stress response kinase PKR is fine-tuned by dynamic clustering. Journal of Cell Biology 221: e202111100

Zhu PJ, Khatiwada S, Cui Y, Reineke LC, Dooling SW, Kim JJ, Li W, Walter P & Costa-Mattioli M (2019) Activation of the ISR mediates the behavioral and neurophysiological abnormalities in Down syndrome. Science 366: 843–849

